# Overexpression of Mig-6 in Limb Mesenchyme Leads to Accelerated Osteoarthritis in Mice

**DOI:** 10.1101/871350

**Authors:** Melina Rodrigues Bellini, Michael Andrew Pest, Jae-Wook Jeong, Frank Beier

**Affiliations:** Department of Physiology and Pharmacology, Western University, London, ON, Canada; Western University Bone and Joint Institute, London, ON, Canada; Children’s Health Research Institute, London, ON, Canada; Department of Obstetrics, Gynecology and Reproductive Biology, Michigan State University College of Human Medicine, Grand Rapids, Michigan

## Abstract

**Background:** Mitogen-inducible gene 6 (Mig-6) is a tumour suppressor gene that is also associated with the development of osteoarthritis (OA)-like disorder. Recent evidence from our lab and others showed that cartilage-specific Mig-6 knockout (KO) mice develop chondro-osseous nodules, along with increased articular cartilage thickness and enhanced EGFR signaling in the articular cartilage. Here, we evaluate the phenotype of mice with skeletal-specific overexpression of Mig-6.

**Methods:** Synovial joint tissues of the knee were assessed in 12 and 36 weeks-old skeleton-specific *Mig-6* overexpressing (*Mig-6^over/over^*) and control animals using histological stains, immunohistochemistry, semi-quantitative OARSI scoring, and microCT for skeletal morphometry. Measurement of articular cartilage and subchondral bone thickness were also performed using histomorphometry.

**Results:** Our results show only subtle developmental effects of Mig-6 overexpression. However, male *Mig-6^over/over^* mice show accelerated cartilage degeneration at 36 weeks of age, in both medial and lateral compartments of the knee. Immunohistochemistry for SOX9 and PRG4 showed decreased staining in *Mig-6^over/over^* mice relative to controls, providing potential molecular mechanisms for the observed effects.

**Conclusion:** Overexpression of *Mig-6* in articular cartilage causes no major developmental phenotype but results in accelerated development of OA during aging. These data demonstrate that precise regulation of the Mig-6/EGFR pathway is critical for joint homeostasis.

## INTRODUCTION

Osteoarthritis (OA) is a failure of joint homeostasis and results in the whole-joint tissue degeneration (1). In fact, OA is a multifactorial disease affecting 630 million individuals worldwide, and the economic impact of OA treatment is estimated at 190 billion dollars in direct and indirect health care costs in North America annually (2,3). OA patients experience limits in daily activities and often suffer from co-morbidities including mental health disorders (4). Treatment for pain and inflammation (analgesics, non-steroidal anti-inflammatory drugs (NSAIDS) and targeted physiotherapy (5) are commonly used to address patients’ symptoms, but no effective pharmacological therapy is currently available to delay disease progression. Future directions for effective OA management rely on better understanding of joint physiology and pathophysiological mechanism to develop disease-modifying therapies for OA patients.

Risk factors including aging, genetics, obesity, and trauma contribute to the dysfunction of joint structures in OA. During the early stages of OA, alteration in chondrocyte physiology including cluster formation and changes in the composition of extracellular matrix (ECM) lead to altered cartilage function (6–8). Gradual degeneration of the articular cartilage, subchondral bone sclerosis, osteophyte development, and synovial inflammation/hyperplasia all contribute to joint degeneration in OA (9–11). Expression of matrix metalloproteinases (MMPs) (i.e., MMP-1 and MMP-13) and aggrecanases (disintegrin and metalloproteinase with a thrombospondin type 1 motif (ADAMTS) (i.e., ADAMTS 1,4,5) is up-regulated in response to inflammatory factors and other signals (12–14). Importantly, the tissues of the whole joint work together to maintain joint homeostasis. Therefore, failure in one joint structure might lead to failure of the whole organ, such as the knee joint (15).

Over the past two decades, epidermal growth factor receptor (EGFR) signaling has been studied in several stages of cartilage development and homeostasis. These studies demonstrate both degenerative and protective roles of this pathway (16), with potential therapeutic implications for OA (17–26). EGFR signaling modulates many canonical signaling pathways including MEK/ERK that have been implicated in cellular proliferation and growth in cartilage and bone, as well as Jun N-terminal kinases (JNKs), PLC-PKC signaling and others (24,27,28). Mitogen inducible gene 6 (Mig-6) is well-known as a negative regulator for EGFR signaling (29). Two different mouse strains with global deletion of *Mig-6* demonstrated bone erosion and spontaneous development of OA-like phenotypes (30,31). Cartilage-specific Mig-6 KO mice display normal early bone development, but show anabolic buildup of articular cartilage, and formation of chondro-osseous nodules at 12 and 36 weeks of age (32). Another study using limb mesenchyme-specific deletion of *Mig-6* in mice (using the *Prx1-cre* driver line) demonstrated similar phenotypes as those observed in cartilage-specific knockout mice (33). Our laboratory has shown that cartilage-specific *Mig-6* overexpression in mice results in no major developmental abnormalities in articular cartilage, however, during aging (12 and 18 months) *Mig-6^over/over^* mice show accelerated cartilage degeneration (34). To evaluate the contribution of Mig-6 in multiple joint tissues to joint homeostasis and OA pathogenesis, we used *Prx1* promoter-driven Cre recombinase to selectively overexpress *Mig-6* in all mesenchymal limb tissues in mice.

## Materials and Methods

### Animals

All animals and procedures were approved by the Council for Animal Care (CCAC) at Western University-Canada (Animal use permit:2015-031). *Mig-6* overexpression animals with the overexpression targeted to the Rosa26 locus (35) were backcrossed for 10 generations into a C57Bl/6 background. In these mice, transcription of *Mig-6* is under the control of a ubiquitously expressed chicken beta actin-cytomegalovirus hybrid (CAGGS) promoter, but blocked by a “Stop Cassette” flanked by LoxP sites (LSL) (35). *Mig-6* overexpression mice were bred to mice carrying the Cre recombinase gene under the control of the Prx1-Cre transgene (36) to induce recombination and removal of the Stop Cassette specifically in early limb bud mesenchyme. Animals with overexpression of Mig-6 from both alleles are termed *Mig-6^over/over^* (*Mig-6^over/over^Prx1-Cre^+/-^*), while control mice are identical but without the Cre gene (denoted “control” for simplicity). Mice were group housed (2 or 4 mice per cage of littermate matched control and overexpression animals), on a standard 12 hour light/dark cycle, and with free access to mouse chow and water. Genotyping and assessment of genomic recombination was performed on DNA samples from ear tissue from mice surviving to at least 21 days of age. Standard polymerase chain reaction (PCR) was performed using primer set P1 and P2 can amplify a 300 bp fragment from the wild-type allele, whereas P1 and P3 can amplify a 450 bp fragment from the targeted ROSA26 locus allele (35).

### Histologic Assessment

The knee joints of mice were dissected and fixed in 4% paraformaldehyde in phosphate buffered saline (PBS, pH 7.0) for 24 hours at room temperature. The intact joints were then decalcified in 5% ethylenediaminetetraacetic acid (EDTA) in phosphate buffered saline (PBS), pH 7.0 for 10 – 12 days at room temperature. All joints were processed and embedded in paraffin in sagittal or frontal orientation, with serial sections taken at a thickness of 5 μm. Sections were stained with Toluidine Blue (0.04% toluidine blue in 0.2M acetate buffer, pH 4.0, for 10 minutes) for glycosaminoglycan content and general evaluation of articular cartilage.

Immunohistochemistry was performed on frontal sections of paraffin embedded knee joints as previously described (32,37). Primary antibodies against SOX9 (R&D Systems, AF3075), MMP13 (Protein Tech, Chicago, IL, USA, 18165-1-AP), and lubricin (Abcam, ab28484) were used and slides without primary antibody were used as control. Sections were incubated with primary antibody overnight at 4°C. After washing, sections were incubated with horseradish peroxidase (HRP)-conjugated donkey anti-goat or goat anti-rabbit secondary antibody (R&D system and Santa Cruz), before incubation with diaminobenzidine substrate as a chromogen (Dako, Canada). Finally, sections were counterstained with 0.5% methyl green (Sigma) and dehydrated in graded series of 70-100% ethanol in water, followed by 100% xylene, and mounted using xylene-based mounting media. All images were taken using a Leica DM1000 microscope with attached Leica DFC295 digital camera.

### Histologic evaluation of articular cartilage and histopathology scoring

Articular cartilage thickness was determined from toluidine blue-stained frontal sections of knee joints by a blinded observer with regard to the tissue source. ImageJ Software (v.1.51) (38) was used to measure the cartilage thickness separately for the non-calcified articular cartilage (measured from the superficial tangential zone to the tidemark) and the calcified articular cartilage (measured from the subchondral bone to the tidemark) across three evenly spaced points from all four quadrants of the joint (medial/lateral tibia and femur), in 4 sections spanning at least 500 μm. For OARSI scoring, Toluidine blue-stained sections were evaluated by one to two blinded observers (MB, MAP) on the four quadrants of the knee: lateral femoral condyle (LFC), lateral tibial plateau (LTP), medial femoral condyle (MFC), and medial tibial plateau (MTP), according to the Osteoarthritis Research Society International (OARSI) histopathologic scale (39). Subchondral bone area from the tibial plateau was traced by one observer (MB) using the Osteomeasure analysis software (OsteoMetrics, Decatur, GA, USA) for histomorphometry measurements using three sections spanning at least 500 μm from each animal.

### Visualization of collagen fiber content

In order to analyze the collagen fibril content and network, Picrosirius Red Staining (0.1% Sirius red in saturated picric acid solution for 60 minutes, with 0.5% acetic acid washes) was performed (32). Stains were imaged under polarized light microscopy to visualize the organization and size of collagen fibrils. Light intensity and tissue angle (45°) relative to polarizing filter (Leica no. 11505087) and analyzer (Leica no. 11555045) were kept identical between samples as per (32).

### Micro-Computerized Tomography (μCT)

Mice were euthanized and imaged using General Electric (GE) SpeCZT microCT machine (40) at a resolution of 50μm/voxel or I00um/voxel in 12 and 36 week-old control and *Mig-6^over/over^* male and female mice. GE Healthcare MicroView software (v2.2) was used to generate 2D maximum intensity projection and 3D isosurface images to evaluate skeletal morphology (32,41). MicroView was used to create a line measurement tool in order to calculate the bone lengths; femurs lengths were calculated from the proximal point of the greater trochanter to the base of the lateral femoral condyle. Tibiae lengths were measured from the midpoint medial plateau to the medial malleolus. Humerus lengths were measured from the midpoint of the greater tubercle to the center of the olecranon fossa.

### Statistical Analysis

All statistical analyses were performed using GraphPad Prism (v6.0). Differences between two groups were evaluated using Student’s *t*-test, and Two-Way ANOVA was used to compare 4 groups followed by a Bonferroni multiple comparisons test. All *n* values represent the number of mice used in each group/genotyping.

## RESULTS

### Overexpression of Mig-6 has minor effects on body weight during development

Mice with alleles for conditional overexpression of Mig-6 (35) were bred to mice expressing Cre recombinase under control of the Prx1 promoter, which is active in the mesenchyme of developing limb buds. Homozygote mice overexpressing Mig-6 in mesenchymal limb tissue from both Rosa26 alleles are referred to as *Mig-6^over/over^* from here on. Control mice do not express Cre recombinase. Overexpressing mice were obtained at the expected Mendelian ratios (data not shown). Animal weights were significantly lower at 7, 12, and 13 weeks after birth in male mutant mice compared to control mice (Fig. 1A), while female *Mig-6^over/over^* mice had similar weights as control mice (Fig. 1B). However, at 36 weeks of age mice there were no differences in weights of either male nor female mutant mice compared to the control group (Fig. 1 C, D).

**Figure 1:**
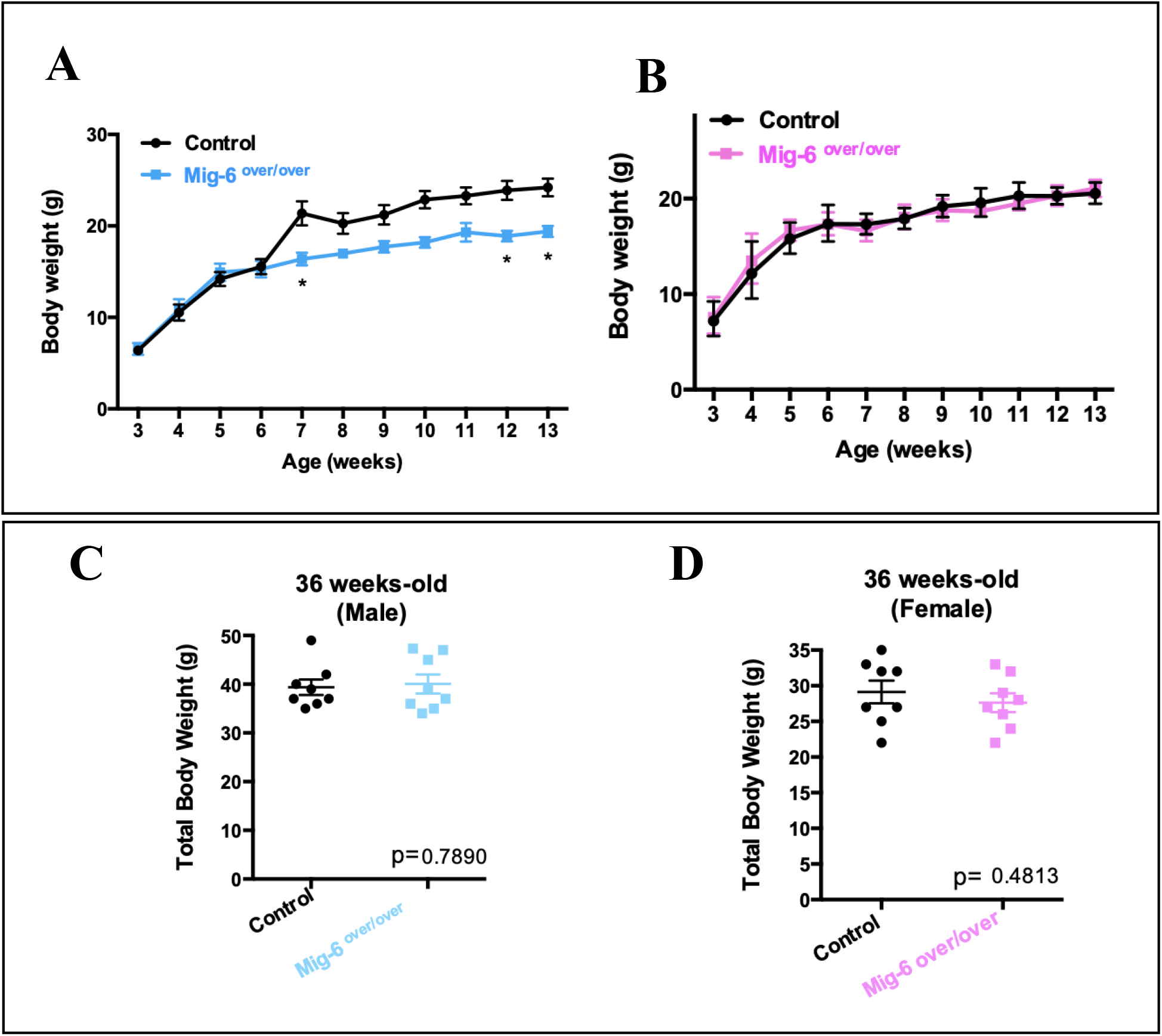
Total body weight of male and female control and *Mig-6^over/over^* mice. Body weight of male Mig-6 overexpressing mice did is significantly lower than control at 7w, 12w and 13w of age **(A)**. Female Mig-6 overexpression mice did not show any statistically significant differences compared to control **(B)**. Two-Way ANOVA was used with Bonferroni post hoc analysis (n=5/genotyping). Data are presented with mean and error ± SEM (P<0.05). Total body weights of 36 week-old male **(C)** and female **(D)** *Mig-6^over/over^* mice and controls did not show any statistically significant differences. Individual data points are presented with mean ± SEM (P<0.05). Data were analyzed by two tailed student t-tests from 8-10 mice per group (age/genotyping).

### Mig-6 overexpressing mice show no differences in bone length

Micro computed tomography (microCT) was used to investigate skeletal morphology and bone length. Whole body microCT scans of *Mig-6^over/over^* male mice and their controls were taken post-mortem at 12 and 36 weeks of age to generate 3D isosurface reconstructions of 50μm/voxel uCT scans, in order to measure long bones lengths (femurs, humeri, and tibiae) in GE MicroView v2.2 software. Mutant male mice at 12 and 36 weeks did not show any difference in bone length compared to controls (Fig. 2A-B). Moreover, no differences in gross skeletal morphology were detected (Fig. 2C).

**Figure 2:**
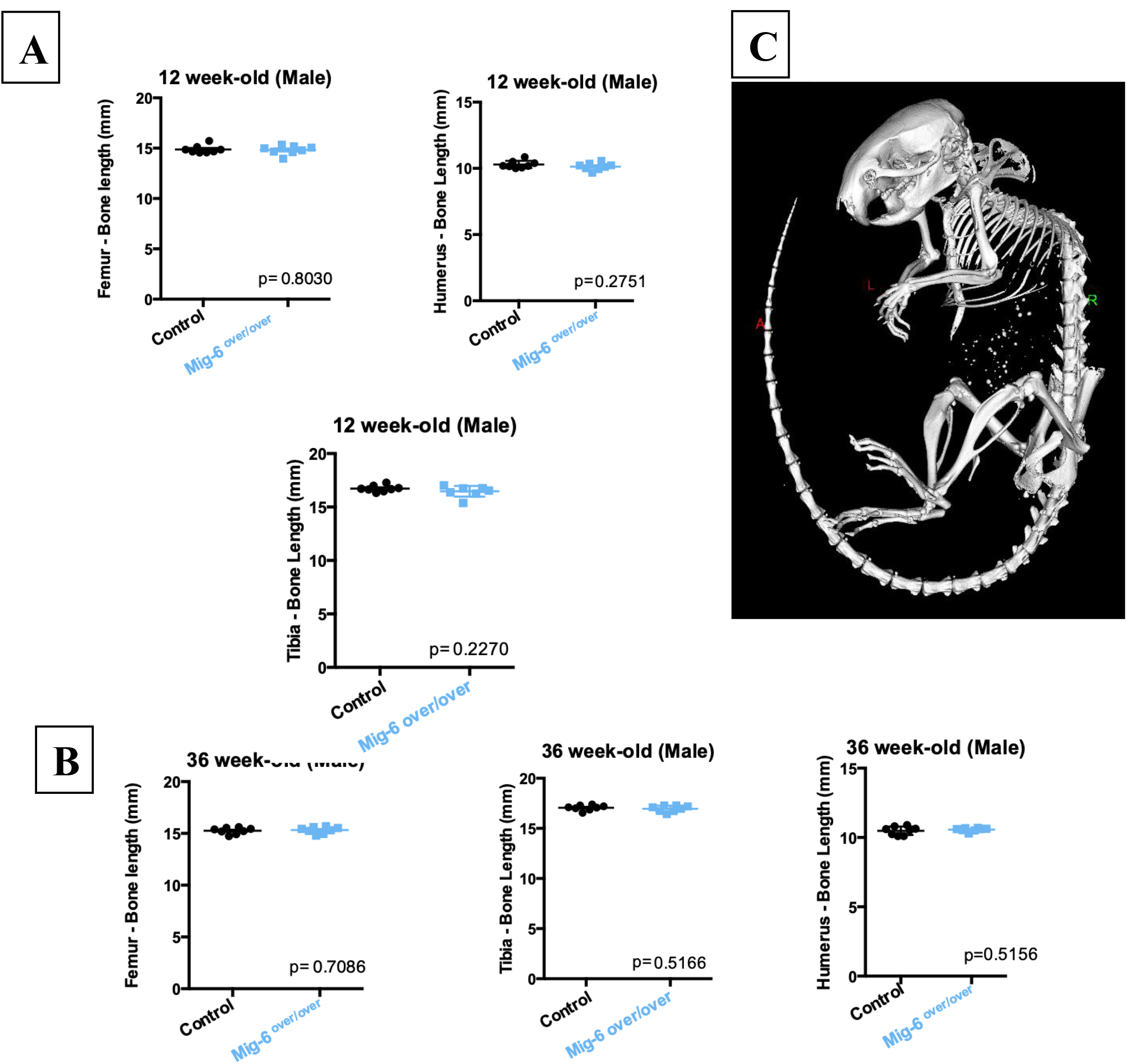
Mig-6 overexpression does not affect bone length. The lengths of right femora, tibiae and humeri were measured on microCT scans of mice at 12 **(A)** and 36 **(B)** weeks of age using GE MicroView software. There were no statistically significant differences in any bones at either age. Individual data points are presented with mean ± SEM (P<0.05). Data were analyzed by two tailed student t-tests from 8 mice per group (age/gender). **(C)** shows a representative 3D isosurface reconstruction of a 100μm/voxel μCT scan.

### Specific overexpression of Mig-6 in limbs display healthy articular cartilage at skeletal maturity

Histologically analysis of knee sections was performed on 12 week-old mutant and control male mice using toluidine blue stained paraffin frontal knee sections (Fig. 3A-B). No major differences in tissue architecture were seen between genotypes. However, the thickness of the calcified articular cartilage in the medial femoral condyle (MFC) and medial tibial plateau (MTP) of male *Mig-6^over/over^* mice was statistically significant lower than in controls. Uncalcified cartilage did not show any differences between genotypes.

**Figure 3:**
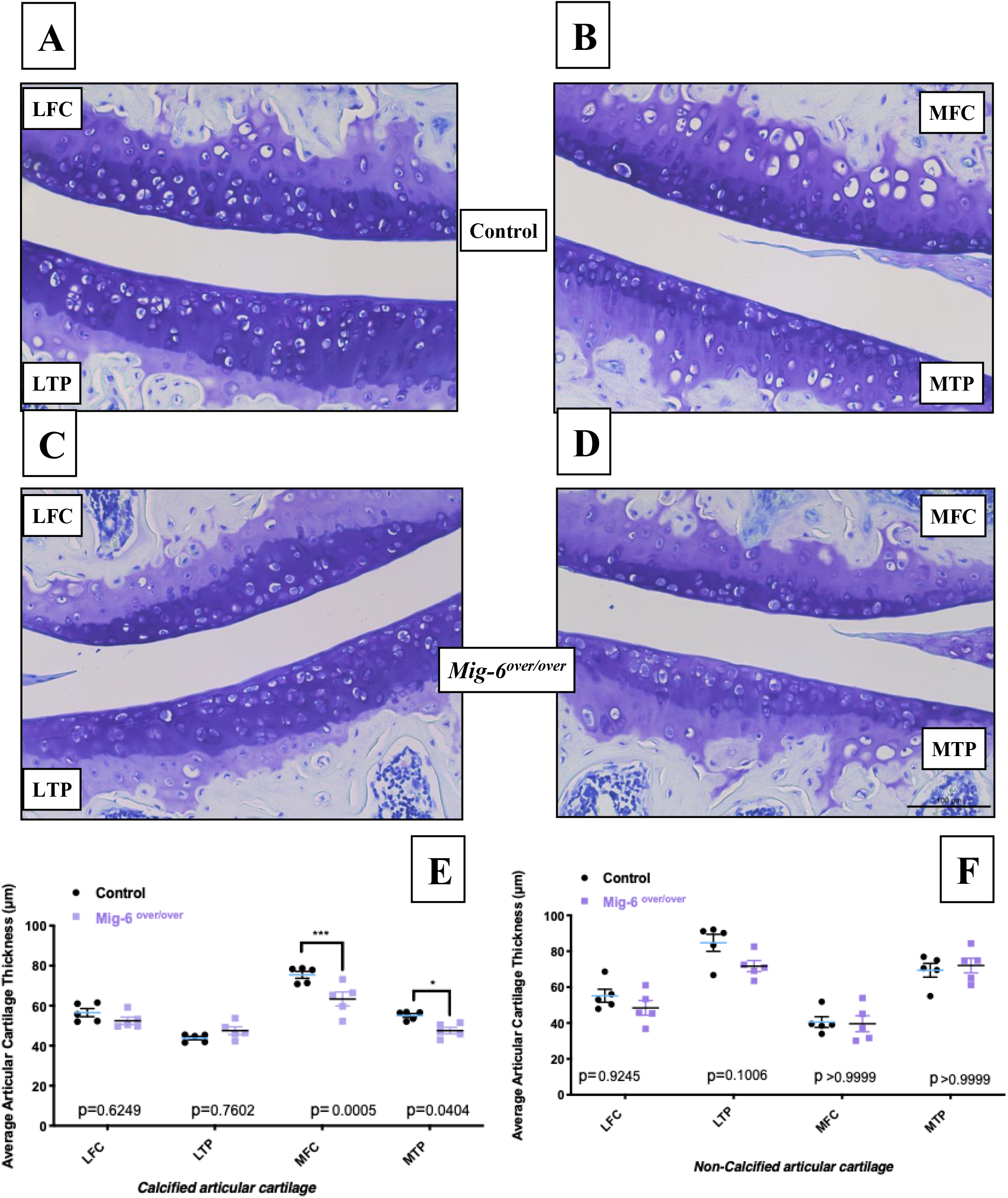
12 week-old *Mig-6^over/over^* male mice show healthy articular cartilage. Representative (n=5) toluidine blue-stained frontal sections of knee joints from 12-week-old control **(A,B)** and *Mig-6^over/over^* **(C,D)** mice showed no apparent damage. Mig-6 overexpressing mice did show statistically significant differences in thickness of the calcified articular cartilage on the medial femoral condyle (MFC) and medial tibial plateau (MTP) **(E)** when compared to controls. However, no statistically significant differences were seen in the non-calcified articular cartilage **(F).** The lateral femoral condyle (LFC) and lateral tibial plateau (LTP) did not show any significant differences. Individual data points are presented with mean ± SEM. Data were analyzed by two-way ANOVA (95% CI) with Bonferroni post-hoc test. Scale bar = 100μm.

### Mig-6 overexpressing male mice display articular cartilage damage at 36 weeks of age

We evaluated the knee joints of 36 weeks-old control and *Mig-6^over/over^* male mice using toluidine blue staining and OARSI grading method (39). At this age, control mice exhibited little to no damage of articular cartilage (Fig. 4A). Conversely, three of seven *Mig-6^over/over^* mice exhibited cartilage damage and erosion with significantly elevated scores in the medial compartment of the knee. Moreover, all seven *Mig-6^over/over^* mice had OA in the lateral compartment of the knee (Fig. 4B), with fibrillation and fissure formation. Furthermore, two of six *Mig-6^over/over^* female mice showed mild cartilage degeneration of the medial compartment (Fig. 5B), in contrast to the control group where no cartilage damage was observed (Fig. 5A).

**Figure 4:**
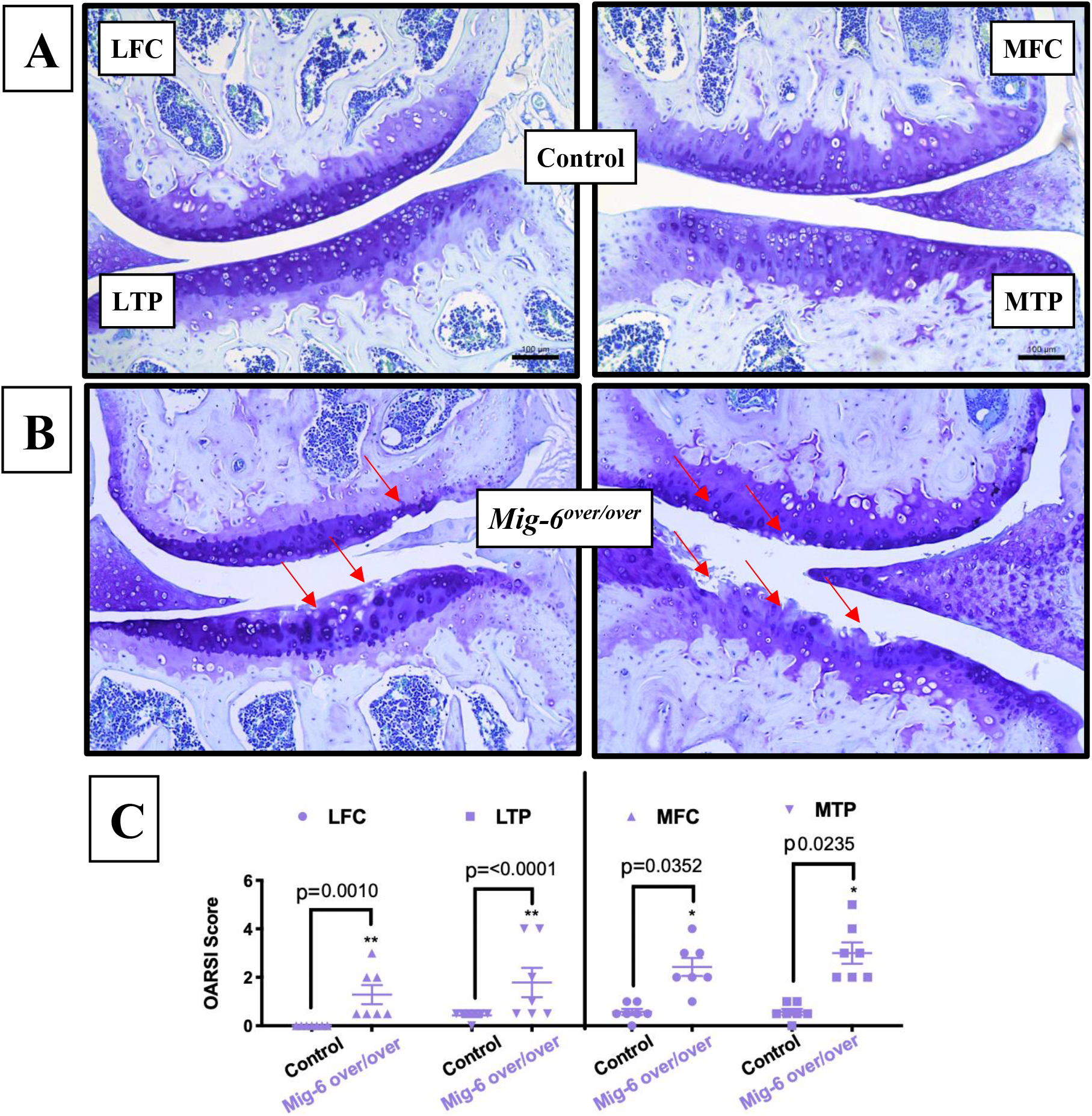
Cartilage damage in knee joints of 36 week-old male Mig-6 overexpressing mice. Toluidine blue staining demonstrated healthy knee joints and articular cartilage in all 36 week-old male control mice **(A)**, while many Mig-6-overexpressing mice showed clear damage to the articular surface **(B)**. OARSI histopathology scoring demonstrated that cartilage degeneration scores significantly increased in the MFC, MTP, LFC and LTP of Mig-6 overexpressing mice. **(C)** Data were analyzed by two-way ANOVA with Bonferroni’s multiple comparisons test. Individual data points are presented with mean ± SEM. All scale bars =100 μm. N = 7 mice/group.

**Figure 5:**
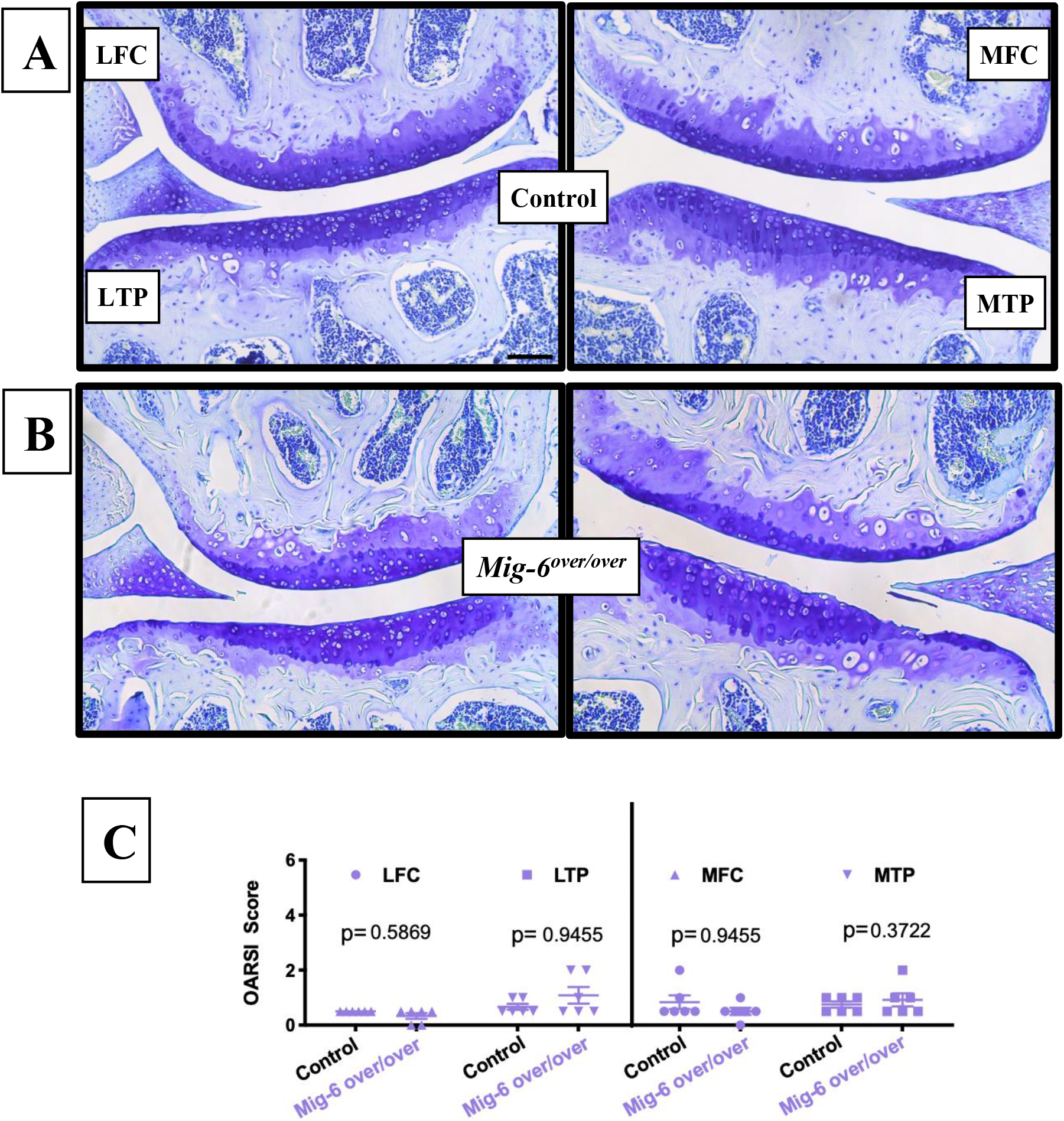
Minor damage in articular cartilage of 36 week-old female Mig-6 overexpressing mice. **(A)** Paraffin sections of knee joints from 36 week-old female control (A) and Mig-6 overexpressing **(B)** mice demonstrated healthy joints in controls and minor cartilage damage in some mutant mice, which was confirmed by OARSI histopathology scoring **(C).** Data were analyzed by two-way ANOVA with Bonferroni’s multiple comparisons test. Individual data points are presented with mean ± SEM. All scale bars =100 μm. N = 7 mice/group.

### Specific overexpression of Mig-6 results in normal bone area

Bone structural alteration is related to knee osteoarthritis as an adaptive response to the loading distribution across joints (42). Measurement of the subchondral bone area from *Mig-6^over/over^* and controls male mice at 36 weeks-old across the entire joint did not reveal any significant differences between genotypes (Fig. 6A-B). Specific measurements of the lateral and medial tibia plateau did not show any significant differences either (data not shown).

**Figure 6:**
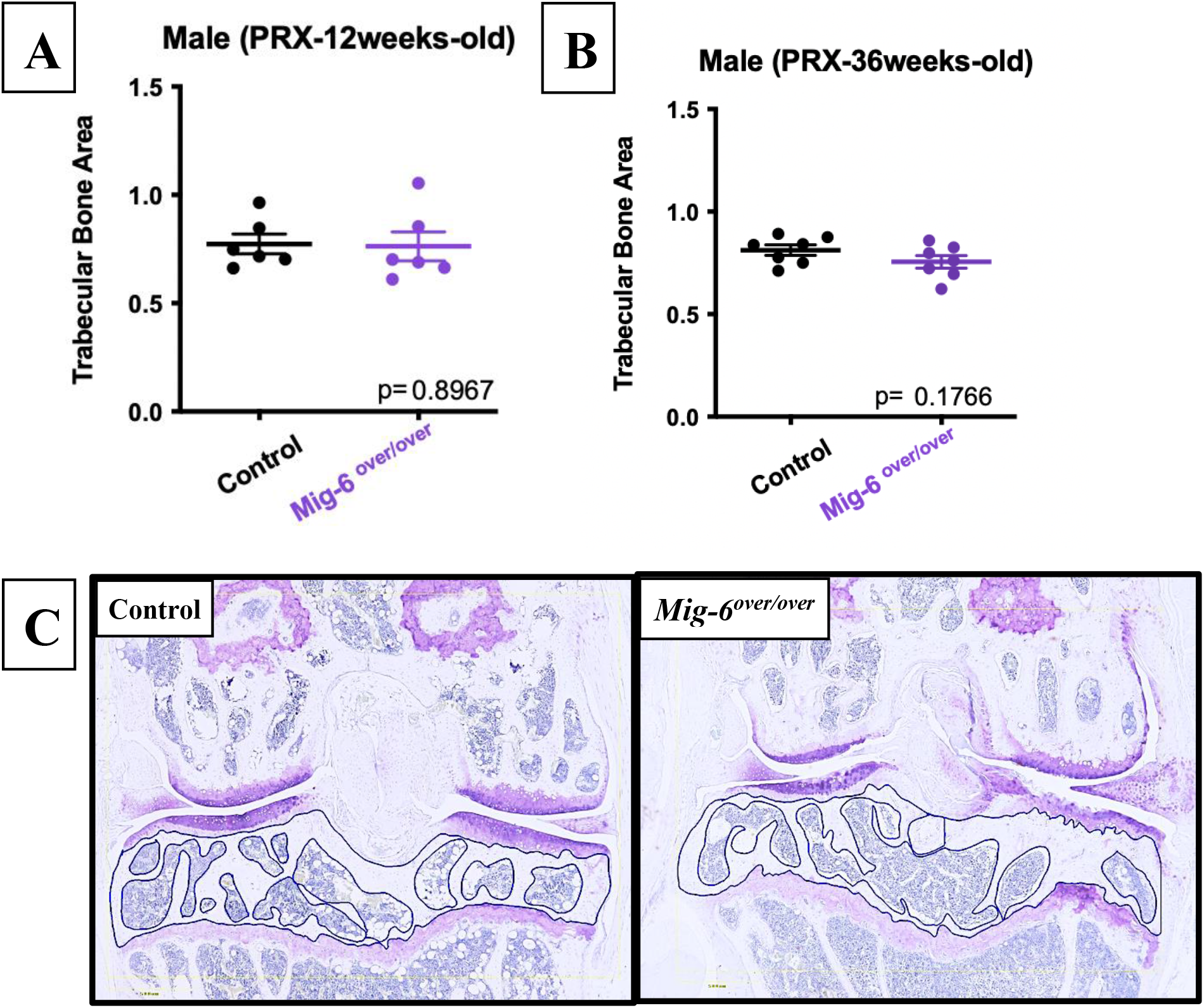
No differences in the subchondral bone area upon overexpression of Mig-6. The subchondral bone area from 12 week-old male control and Mig-6 overexpressing **(A)** or 36 week-old male control and Mig-6 overexpressing **(B)** mice are shown. Representative images of the subchondral area selected using the OsteoMeasure bone histomorphometry system are shown in **(C)**. Individual data points are presented with mean ± SEM. Data were analyzed by one observer (MB). All scale bars =100 μm. N = 6-7 mice/group.

### Mig-6 overexpressing mice display altered collagen fiber organization in articular cartilage

Frontal sections from 36 weeks-old male mice were stained with Picrosirius red to visualize the collagen network under polarized light microscope. In the control male mice, the collagen fibers in the articular cartilage exhibit greenish/yellow birefringence in the superficial and transitional zones, resulting from thin collagen fibers in these regions. In the deep and calcified cartilage, and in bone, red birefringent fibers are visualized, indicating larger fiber diameter in these regions. The articular cartilage of *Mig-6^over/over^* showed fewer green collagen fibers in the medial compartment of the knee, indicating a loss of normal collagen fibers (Fig. 7A-B).

**Figure 7:**
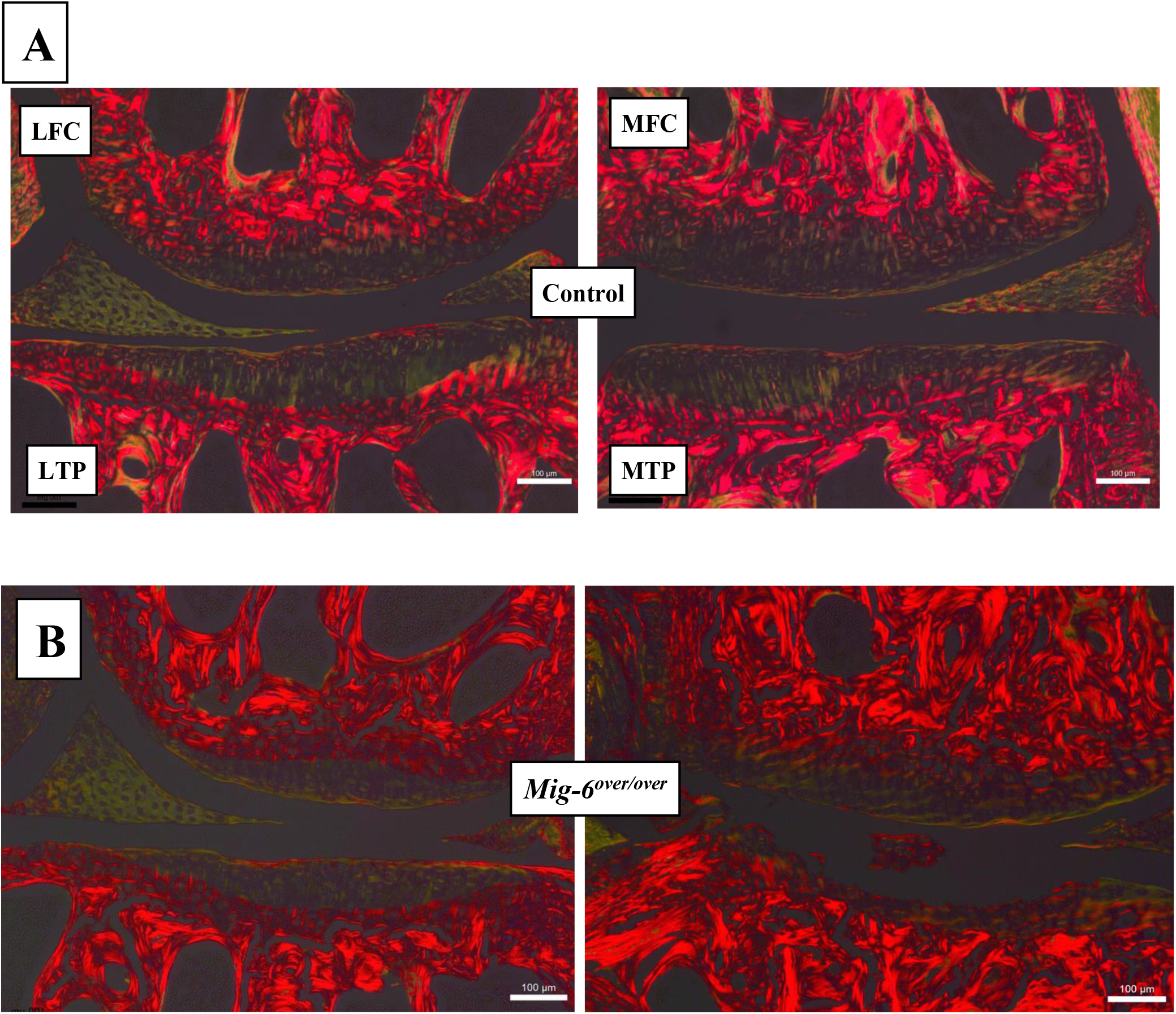
Picrosirius Red Staining of control and Mig-6 overexpressing mice. Representative paraffin sections of the medial and lateral compartment in 36 week-old male control **(A)** and Mig-6 overexpressing mice **(B)** were stained with picrosirius red (fibrillar collagen) and analyzed under polarized light to evaluate the collagen tissue organization and orientation in the articular cartilage. Cartilage in the medial compartment of *Mig-6^over/over^* mice shows reduced collagen staining. N=5 mice/group; LFC = lateral femoral condyle, LTP = lateral tibial plateau, MFC = medial femoral condyle and MTP = medial tibial plateau. Scale bar = 100μm.

### Overexpression of *Mig-6* decreases Sox9 expression

Studies have shown that expression of the transcription factor SRY (sex determining region Y)-box 9 [SOX9] was increased in articular cartilage upon both *Prx1-Cre 1* (43) and *Col2-Cre*-driven deletion of *Mig-6* (32). SOX9 is an essential regulator of chondrogenesis and the maintenance of a chondrocyte-like phenotype (44). Frontal sections of paraffin embedded knees from 12 and 36 week-old male mice were used for SOX9 immunostaining. In 12 weeks-old control mice, nuclear SOX9 was abundantly present in the articular cartilage of the knee joints in all four quadrants. In contrast, *Mig-6^over/over^* mice appear to have fewer cells staining positive in both lateral and medial compartments (Fig. 8A,B). 36 week-old *Mig-6^over/over^* mice showed a further reduction in SOX9 immunostaining in the lateral quadrant, while the loss of cartilage in the medial compartment led to an absence of SOX9 staining (Fig. 9A,B). For both ages, negative controls did not show staining in chondrocytes (data not shown).

**Figure 8:**
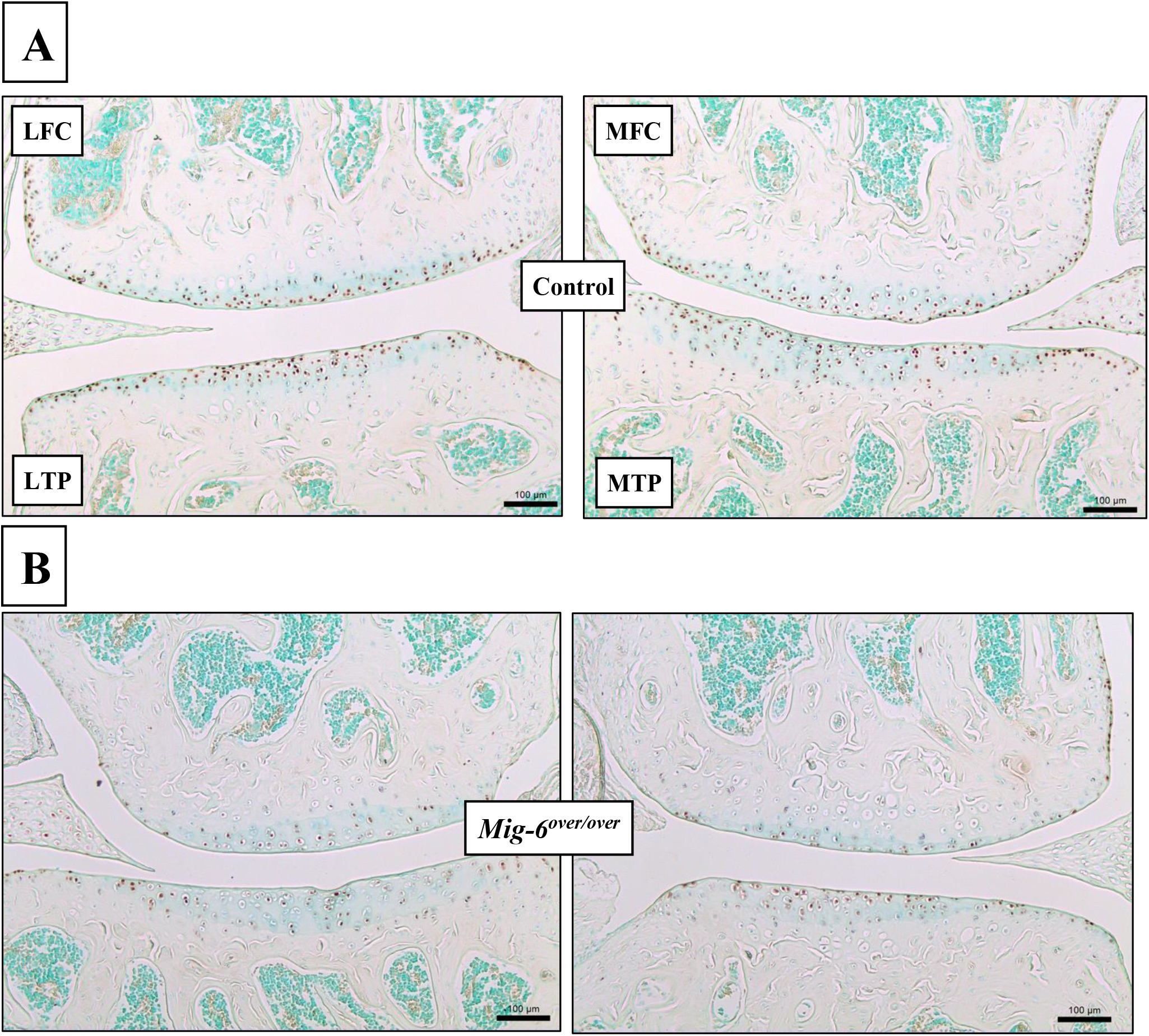
Lower numbers of SOX9-positive cells in 12-week old male Mig-6 overexpressing mice. Representative SOX9 immunostaining in knee joints of 12 week-old male control **(A)** or Mig-6 overexpressing **(B)** mice (n=5 mice/group). Overexpressing mice showed reduced numbers of positive cells in the medial and lateral compartments. LFC = lateral femoral condyle, LTP = lateral tibial plateau, MFC = medial femoral condyle and MTP = medial tibial plateau. Scale bar = 100μm.

**Figure 9:**
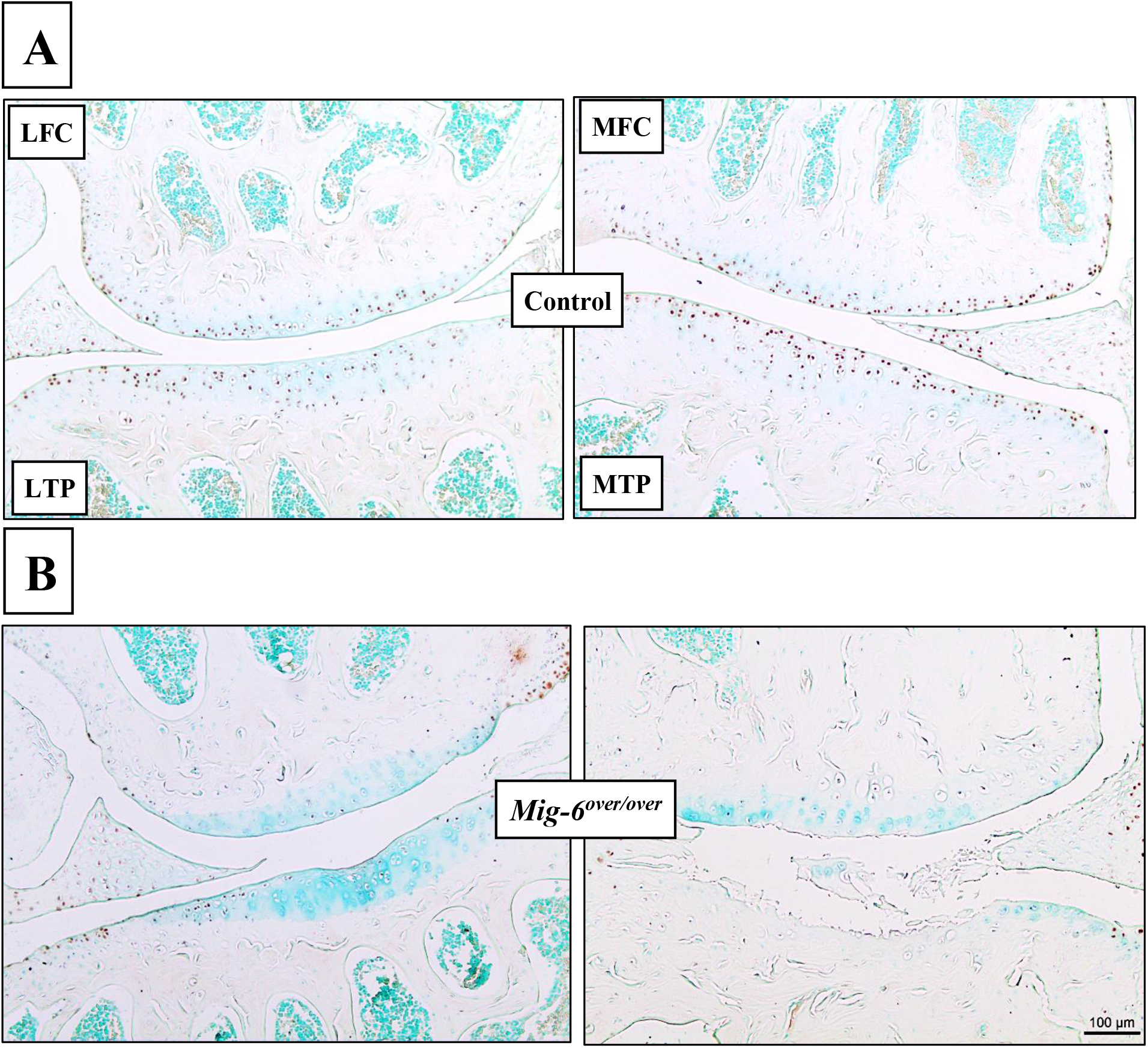
Lower numbers of SOX9-positive cells in 36-week old male Mig-6 overexpressing mice. Representative SOX9 immunostaining in knee joints of 36 week-old male control **(A)** or Mig-6 overexpressing **(B)** mice (n=5 mice/group). Overexpressing mice showed reduced numbers of positive cells in the medial and lateral compartments. LFC = lateral femoral condyle, LTP = lateral tibial plateau, MFC = medial femoral condyle and MTP = medial tibial plateau. Scale bar = 100μm.

### Lubricin/PGR4 is slightly decreased upon Mig-6 overexpression

Lubricin/proteoglycan 4 plays an important role as joint boundary lubricant and is produced by synoviocytes as well as superficial zone chondrocytes (45,46). In 12 week-old *Mig-6^over/over^* mice, lubricin was observed in superficial zone (SZ) and middle zone (MZ) chondrocytes, in a similar pattern as control mice, although intensity appeared reduced in overexpressors (Fig. 10 A-C). 36 weeks-old control male mice show lubricin immunostaining in the SZ and MZ, however, less lubricin immunostaining is present in the SZ of the medial side of *Mig-6^over/over^* mice. Negative controls did not show staining in chondrocytes or articular cartilage at either age (Fig. 11 A-C).

**Figure 10:**
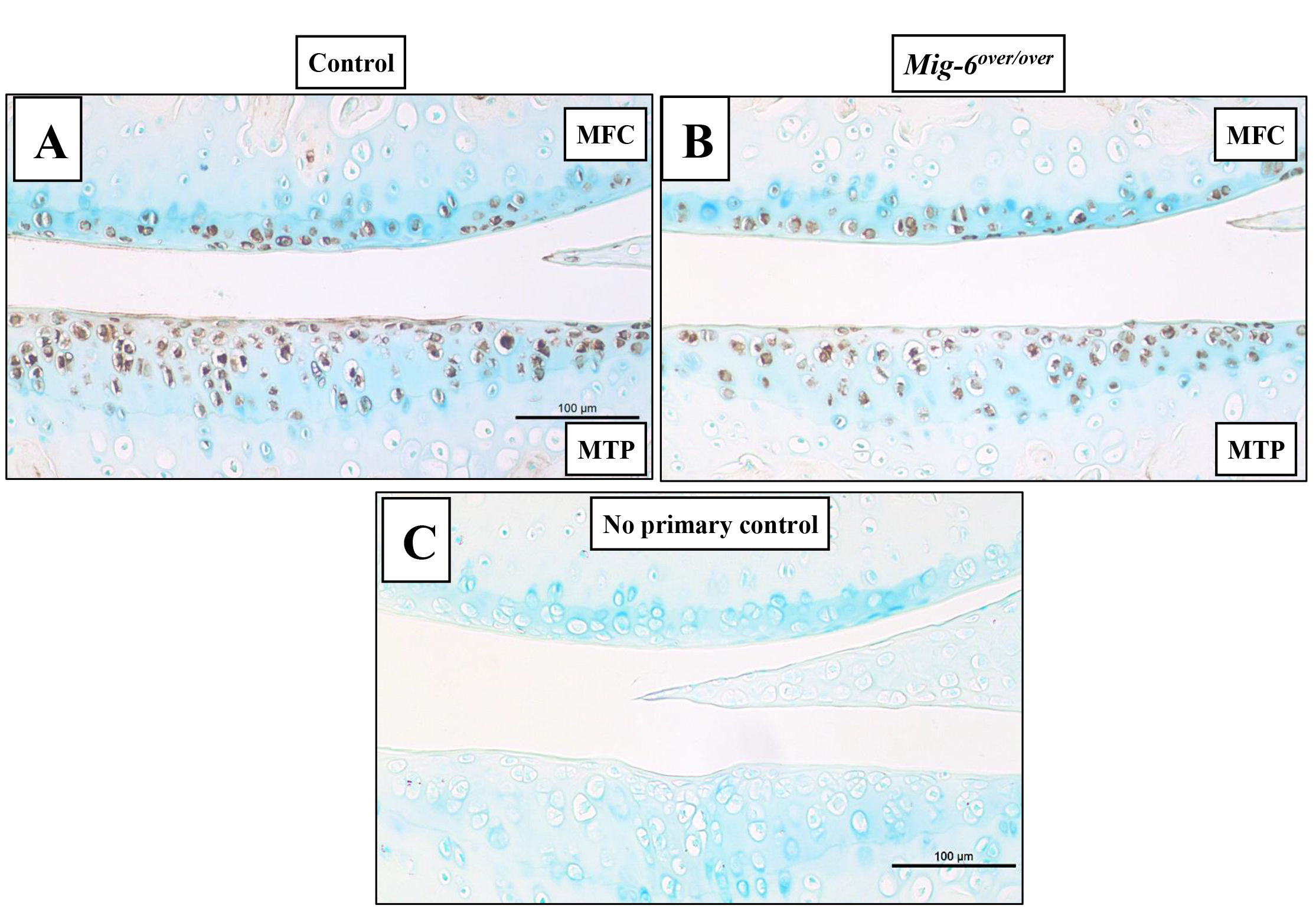
Lubricin immunostaining is slightly decreased in the articular cartilage of Mig-6 overexpressing mice at 12 weeks of age. Immunostaining of sections of the knee joint indicate the presence of Lubricin (*PRG4*) in the superficial zone chondrocytes of 12 week-old male control mice **(A)**, with apparently reduced staining in Mig-6 overexpressing mice **(B)**. Frontal sections of articular cartilage with no primary antibody as negative control exhibited no staining **(C)**. N=5 mice/genotyping. MFC = medial femoral condyle and MTP = medial tibial plateau. Scale bar = 100μm.

**Figure 11:**
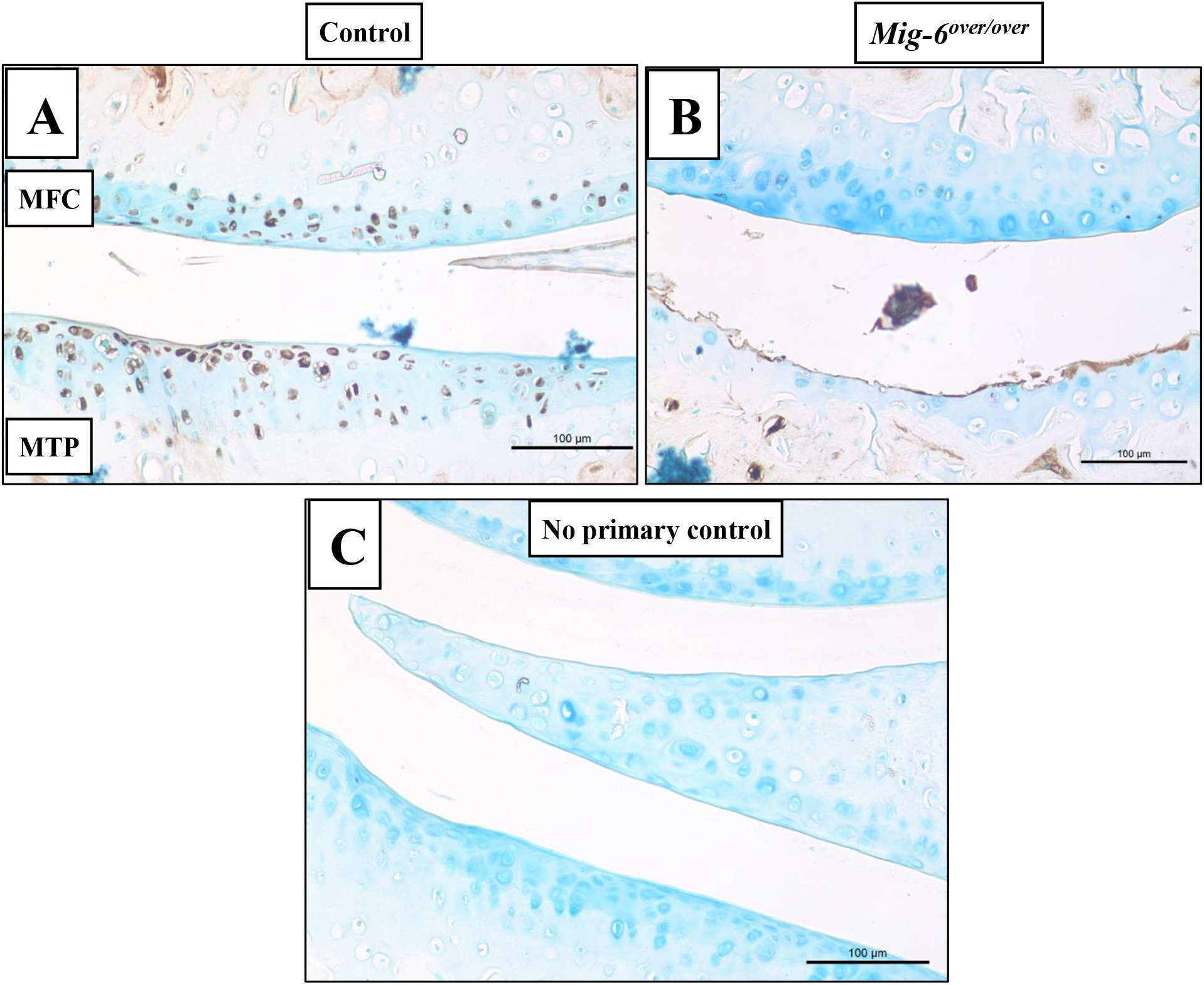
Lubricin immunostaining is decreased in the articular cartilage of Mig-6 overexpressing mice at 36 weeks of age. Immunostaining of sections of the knee joints of 36 week-old male mice indicate the presence of Lubricin (*PRG4*) in the superficial zone of control mice (A), with markedly reduced signal in Mig-6 overexpressing mice **(B)**. Frontal sections of mice articular cartilage with no primary antibody as negative control exhibited no staining **(C)**. N=5 mice/genotyping. MFC = medial femoral condyle and MTP = medial tibial plateau. Scale bar = 100μm.

### MMP13 immunostaining is increased in Mig-6-overexpressing compared to control mice

Previous studies have shown that matrix metalloproteinase (MMP) 13 is the main collagenase associated with type II collagen destruction in OA. Frontal sections of knees from 36 week-old control and *Mig-6^over/over^* male mice were used for MMP13 immunohistochemistry. While some staining was seen in control mice, intensity of staining was increased in areas of damage on the medial side of *Mig-6^over/over^* mice. Negative controls did not show staining in cartilage or subchondral bone (Fig. 12 A,B and C).

**Figure 12:**
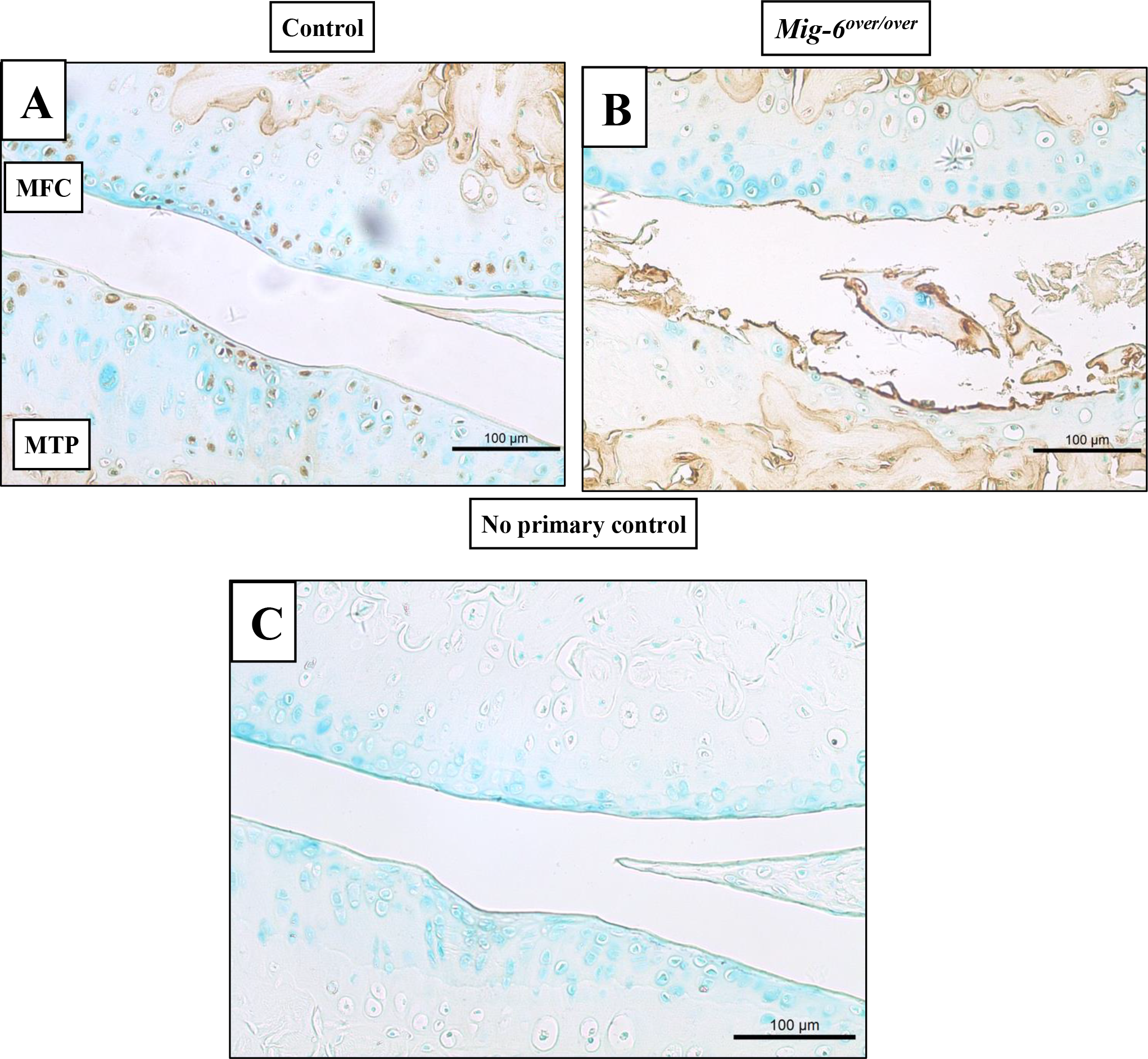
36 week-old Mig-6 overexpressing mice show increased MMP13 staining in cartilage. Representative immunohistochemistry of matrix metalloproteinase 13 (MMP13) in 36 week-old control **(A)** and Mig-6 overexpressing **(B)** mice show increased staining in the degrading cartilage of overexpressors. No primary antibody control is shown in **(C).** N=5 mice/genotyping. LFC = lateral femoral condyle, LTP = lateral tibial plateau, MFC = medial femoral condyle and MTP = medial tibial plateau. Scale bar = 100μm.

## Discussion

Mig-6 has been studied in a variety of human diseases, including cancer and more recently OA progression(16,31–33,47). Many studies, including from our lab, also identified TGFα /EGFR signaling as a regulator of OA progression and cartilage homeostasis (21,24,48). Interestingly, cartilage-specific (Col2-Cre) deletion of *Mig-6* (Mig-6 KO) (32) results in increased proliferation of chondrocytes and a thicker layer of cartilage while skeletal-specific (Prx1-Cre) deletion of *Mig-6* results in transient anabolic buildup of cartilage followed by catabolic events such as cartilage degeneration at 16 weeks of age (33). In fact, global deletion of *Mig-6* in mice results in a complex set of phenotypes, including joint damage at relatively early time-points in a surgical mouse model (31,49,50). Previous research demonstrates that Mig-6 acts as a negative feedback inhibitor of EGFR signaling (51). Thus, Mig-6 has been suggested as a potential tumor suppressor, as a suppressor of EGFR signaling in human carcinomas (35,52–55). Recent work has revealed that overexpression of Mig-6 acts as a negative regulator of EGFR-ERK signalling in mouse uterus (35). In our study, we set out to evaluate the role of *Mig-6* in joint physiology by using skeletal-specific constitutive overexpression of *Mig-6*. In this study, we show no major effects of Mig-6 overexpression on bone length at the ages of 12 or 36 weeks. While male *Mig-6^over/over^* mice did show slightly reduced body weight up to 12 weeks after birth, these differences were no longer present at 36 weeks of age.

Our results show that *Mig-6^over/over^* (Prx1-Cre) male mice developed cartilage lesions at 36 weeks of age, where control mice show healthy cartilage. OARSI scores of *Mig-6^over/over^* mice reveal significantly increased cartilage degeneration compared to control group. Surprisingly, it appeared that cartilage degeneration in *Mig-6^over/over^* mice was not accompanied by any obvious changes in subchondral bone. However, the thickness of the calcified articular cartilage in *Mig-6^over/over^* was significantly decreased at the 12-week time-point, at least in the medial compartment. It is currently unclear whether and how this is related to the subsequent degeneration of articular cartilage in these mice.

SOX9 is a transcription factor that is necessary for the formation of mesenchymal condensations as well as chondrocyte differentiation and proliferation (56,57). Our data suggest a lower number of SOX9-positive cells at the 12-week time point in mutant mice, preceding cartilage damage. The number of SOX9-expressing cells is further reduced in 36 week-old mutant mice, although this is partially due to the loss of cartilage and chondrocytes. In agreement with these data, mice with cartilage*-* or limb mesenchyme-specific deletion of *Mig-6* showed increased expression of SOX9 in the articular cartilage (32,33).

Lubricin/*PRG4* is necessary for joint lubrication and to maintain healthy cartilage (58,59).

Our results suggest a slight decrease in lubricin staining in 12 weeks-old male *Mig-6^over/over^* mice, compared to the control group. We also observed the same trend towards decreased staining in 36 weeks-old male Mig-6 over/over mice. Together, these data suggest that the decreased of SOX9 and lubricin in the articular cartilage could contribute to cartilage degeneration in our mutant mice.

We recently described mice with cartilage-specific (Col2-Cre-driven) overexpression of Mig-6 (34). Despite the differences in recombination patterns conferred by the two different Cre drivers, overall the phenotypes observed upon Mig-6 overexpression are quite similar. Both are characterized by no or only subtle developmental defects, followed by reduced SOX9 and lubricin expression, followed by cartilage degeneration. One unique feature of the Prx1-driven Mig6-overexpression described here is the stronger OA phenotype in the lateral compartment of 36 week-old mutant male mice. Future studies will need to investigate the underlying causes.

While Mig-6 had been identified as a negative regulator of EGFR signaling, it also interacts with a number of other potential candidate proteins that may contribute to the phenotype described here, such as Cdc42 (60), c-Abl (61), and the hepatocyte growth factor receptor c-Met (62). Therefore, additional work is necessary to elucidate the potential role of these proteins in the phenotype presented. In conclusion, in this study using limb mesenchyme-specific *Mig-6* overexpression we show a reduction of SOX9 and PRG4 expression, and accelerated cartilage damage. The data highlights the importance of more studies on the specific role of Mig-6 signaling in joint homeostasis and OA development.

## Acknowledgements

We would like to thank Julia Bowering for performing joint sectioning. We thank all members of the Beier lab for discussions and help. M.B. was supported by a fellowship from CNPq/Brazil. Work in the lab of F.B. is supported by a grant from the Canadian Institutes of Health Research (Grant #332438). F.B. holds the Canada Research Chair in Musculoskeletal Research.

## Notes

The authors have no conflicts of interest to declare.

